# Jamaican fruit bats are susceptible to henipaviruses but rapidly control infection

**DOI:** 10.64898/2026.07.17.739140

**Authors:** Julianne Bullock, Brown Bulloch, Arthur Wickenhagen, Shane Gallogly, Franziska Kaiser, Jonathan E. Schulz, Trenton Bushmaker, Reshma Koolaparambil Mukesh, Brandi N. Williamson, Kailin Hawes, Anthony McBain, Jessica Prado-Smith, Chad S. Clancy, Brian J. Smith, Julia R. Port, Emmie de Wit, Sarah van Tol

## Abstract

Henipaviruses (HNV), Hendra (HeV) and Nipah (NiV) virus, cause severe pulmonary and neurologic disease in humans and other mammals. Bats in the genus *Pteropus* naturally host HeV and NiV, but their feasibility in experimental studies is limited due to their large size, low fertility rate, and unavailability outside of their native range. Understanding bat-henipavirus interactions that regulate shedding and replication could improve mitigation of spillover events and illuminate the factors that differentiate severe and controlled HNV infection. Here, we assessed the suitability of the Jamaican fruit bat (JFB) (*Artibeus jamaicensis*) to model HNV infection *in vitro* and *in vivo*. JFB primary kidney cells were permissive to HNVs, and HeV and NiV antagonized the induction of the innate antiviral response. JFBs were inoculated via the intranasal and oral routes (IN/PO) with HeV or NiV or intravenously (IV) with HeV and monitored for 7 days. Following IN/PO exposure, infection was quenched rapidly and limited HeV RNA was detected in oral swabs and tissues while NiV RNA was found in only one oral swab. HeV IV inoculation resulted in robust, disseminated infection and viral RNA was detected in oral, rectal, and environmental swabs. Overall, these results support that JFBs are susceptible to both viruses, but replication is quenched rapidly *in vivo* following IN/PO exposure. Future studies will optimize the *in vivo* model to leverage the JFB to further our understanding of bat-henipavirus interactions.

**Author Summary:** Henipaviruses (HNV) spillover from pteropid bat species and cause severe diseases in humans. Many gaps limit our understanding of HNV-bat interactions that influence spillover and effective control of infection. Here, we evaluate the Jamaican fruit bat (JFB) as a henipavirus model. We demonstrate that JFB cells are permissive all HNVs evaluated, and that JFBs support Hendra virus replication and shedding without signs of clinical disease.

## Introduction

Henipaviruses (HNV), including Hendra (HeV), Nipah (NiV), and Cedar (CedV) viruses, are negative-sense RNA viruses within the family Paramyxoviridae. In humans, HeV and NiV cause severe clinical manifestations, including acute respiratory distress syndrome and acute encephalitis syndrome (1–7). NiV includes two main strains, NiV-Malaysia (NiV-M) and -Bangladesh (NiV-B). Although NiV-M caused one large outbreak beginning in 1998, NiV-B spillovers into human populations have occurred regularly in India and Bangladesh (8). The two NiV strains combined have caused approximately 700 human cases with a case fatality rate exceeding 70% (9) Although HeV has only infected seven humans (57% case fatality rate) following exposure to infected horses, the virus regularly spills over into Australian horses and causes lethal disease (9).

In contrast to the pathogenic HNVs, CedV is predicted to be apathogenic in humans. HNVs phosphoprotein gene (P) encodes an RNA-editing site that introduces non-encoded guanine (G) residues into viral transcripts (10) This frameshift results in the production of accessory viral proteins V (+1 G) and W (+2 G) which are key innate antiviral antagonists (11–15) and determinants of pathogenesis (16–19). Unlike HeV and NiV, CedV P does not encode an RNA-editing site, thus V and W are not produced (20, 21).

Bats in the family Pteropodidae are the natural HNV hosts. Australia flying foxes (*Pteropus spp.*) are the reservoir for HeV (22–24) and CedV (20), while various pteropid bats in Southeast Asia host NiV (25–30). Although HNV spillovers to humans have only been documented in Australia, Malaysia, Bangladesh, the Philippines, and India, molecular and serological evidence supports that pteropodid species host diverse HNV and HNV-like viruses throughout their natural range (31–34). The disparity between HNV’s geographic distribution and spillover raises questions about the biotic and abiotic determinants of viral shedding from pteropid bats and of resulting disease in spillover hosts.

Unlike humans, HNV infected pteropid bats do not present with overt clinical disease (35–38). Several challenges limit the practicality of pteropid bats as a tractable experimental model. For example, their large body size, slow reproductive rate, and limited availability outside their native range limit the sample sizes for experimental infection studies. These restrictions have resulted in limited data being generated and relatively underpowered studies (35–39). The existing data suggests that within naturally and experimentally infected pteropid bats, HNV RNA is detected most frequently in the kidney and spleen and shed in urine and oral secretions (40–42). No gross pathology is observed at necropsy of experimentally inoculated bats, and only minor histopathologic lesions, typically localized vasculitis, are detected (35).

Most of these published studies have used wild-caught bats (35–37). Although the bats were shown to be negative for HNV-neutralizing antibodies, evidence supports that free-ranging pteropids can serorevert and are susceptible to reinfection (43). The unknown HNV infection history may confound the results of HNV infection studies on wild-caught bats. Overall, these limitations minimize the capacity to evaluate HNV-bat host dynamics experimentally with the natural reservoir.

Here, we evaluated the suitability of the Jamaican fruit bat (JFB), *Artibeus jamaicensis*, to model HNV infection. We show that JFBs are susceptible to HeV, but the infection was self-limiting after intranasal and oral (IN/PO) inoculation. Following HeV intravenous (IV) inoculation, we observe robust infection of the kidney and spleen as well as oral shedding of infectious virus. Despite *in vitro* studies supporting similar susceptibility of JFB primary kidney cells to both HeV and NiV, HeV RNA was detectable in oral swabs and tissue samples at a higher frequency than NiV RNA. Overall, these results support the potential to develop the JFB as a tractable model to evaluate the environmental and biological factors that influence henipavirus susceptibility, pathogenesis, and shedding.

## Results

### Jamaican fruit bat cells are permissive to henipaviruses

To investigate whether JFB cells support henipavirus replication, we infected primary kidney cells (AjKi2) with CedV, HeV, NiV-M, or NiV-B. When inoculated with a low multiplicity of infection (MOI), 0.005, to monitor multiple cycles of replication, AjKi2 cells supported replication of all four HNVs evaluated (Fig 1A). HeV replicated to higher titers than NiV-B at 24- and 72-168 hours post-inoculation (HPI), but the infectious titers did not significantly differ from NiV-M (Fig 1A). Of note, CedV replication peaked at 72 HPI, and infectious virus declined through the end of sample collection (Fig 1A) despite viral RNA levels remaining stable (Fig 1B). After infection with MOI 1.0, HeV replicated to higher titers than NiV-B, 24- and 48 HPI, and NiV-M 48 HPI (Fig 1C), though viral RNA was only significantly lower for NiV-B (Fig 1D). Insufficient stock titers prevent CedV from being evaluated at MOI 1.0.

**Fig 1.**
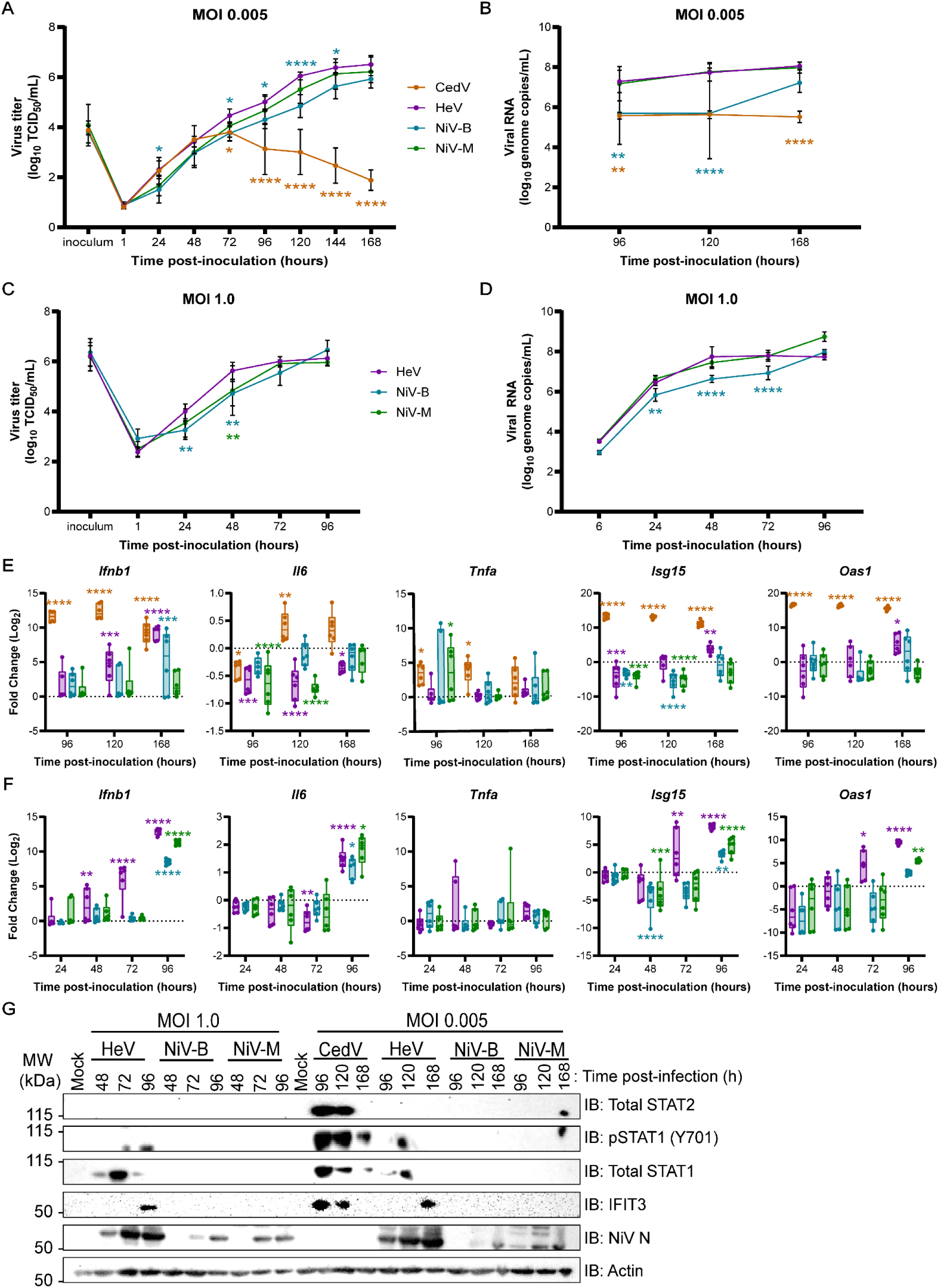
Jamaican fruit bat cells are permissive to henipaviruses. Jamaican fruit bat primary kidney cells were inoculated with Cedar virus (CedV), Hendra virus (HeV), Nipah virus Bangladesh (NiV-B), or NiV Malaysia (NiV-M) at multiplicity of infection (MOI) 0.005 (A-C,G) or 1.0 (D-G) for the indicated times post-inoculation, hours (h). Supernatants were titered to determine the tissue culture infectious dose 50 (TCID50/mL), limit of detection 0.8 TCID50/mL (A, C). Cells were lysed to collect RNA to determine viral load (B, D) and evaluate host gene expression (E, F) using RT-qPCR. Viral loads were determined using a standard curve, and for host gene expression ΔC_T_ values were normalized to the average ΔC_T_ value of mock infected cells to calculate ΔΔC_T_. Protein was collected and immunoblotted to evaluate host and viral proteins (G). Data from three biological replicates (A-F) or one replicate representative of three independent experiments (G) are shown. Data plotted as mean ± S.D. (A-F). Difference in virus titer (A, C) or viral load (B, D) compared to HeV or fold change of each host gene at each time point for infected cells was compared to mock (E, F) using a two-way ANOVA with Dunnett’s multiple test comparison. *P* values for comparisons HeV-inoculated cells (A-D) or to mock cells (E, F) are indicated. **** <0.0001, *** <0.001, ** <0.01, * <0.05.

Cellular RNA and protein samples were collected at all time points to evaluate the host innate antiviral response. As expected, due to the lack of V and W genes, CedV induced robust transcription of *Ifnb1*, pro-inflammatory cytokines, *Il6* and *Tnfa*, and interferon-stimulated genes (ISG), *Isg15* and *Oas1* at either 96-(*Ifnb1*, *Tnfa*, *Isg15*, *Oas1*) or 120 (*Il6)* HPI despite low levels of infectious virus (Fig 1E). In contrast, HeV infection suppressed the transcription of *Ifnb1* and ISGs until 120 HPI and pro-inflammatory cytokines throughout 168 HPI (Fig 1E). NiV-B and NiV-M largely suppressed the innate antiviral response throughout infection, though mild induction of *Ifnb1* transcripts was detected at 168 HPI in NiV-B infected cells (Fig 1E). The same trend was largely observed following inoculation at MOI 1.0. HeV induced transcription of *Ifnb1* (48 HPI) and ISGs (72 HPI), while the same genes were not induced until 96 HPI following NiV-B or NiV-M infection (Fig 1F).

Western blot data correlated with the innate immune gene transcription for both MOIs. As expected, CedV infection induced phosphorylation of STAT1 (pSTAT1) and expression of ISGs (STAT2, STAT1, IFITM3) by 96 HPI and the levels of STAT1 activation and ISG expression decreased by 168 HPI (Fig 1G). For both MOI 0.005 and 1.0, only HeV induced pSTAT1 and expression of ISGs (Fig 1G). Compared to CedV infection, HeV induction of the innate antiviral response was delayed. Overall, these results support that JFB cells are susceptible to henipavirus replication and that HeV and NiVs antagonize JFB innate antiviral response.

### Jamaican fruit bats rapidly control Hendra and Nipah virus after intranasal and oral inoculation without signs of disease

To evaluate susceptibility *in vivo*, JFBs were inoculated with either HeV (n=8) or NiV-M (n=8) via the IN/PO routes and samples were collected to monitor shedding, virus dissemination, and host responses to infection (Fig 2A). Throughout the experiment, bats infected with HeV or NiV did not differ in body temperature (Fig 2B) and weight (Fig 2C) compared to healthy controls. Low levels of HeV RNA were detected in oral (max 2.8 log_10_ copies/mL) and rectal (max 2.2 log_10_ copies/mL) swabs (Fig 2D), with the peak for oral swabs being 2 days post-inoculation (DPI) and rapidly declining, and rectal swab positivity being sporadic. Only one oral swab on day 2 was positive for NiV-M (2.1 log_10_ copies/mL) (Fig 2D). All swab samples, for both HeV and NiV-M were negative for infectious virus. For tissue samples, HeV RNA was detected in spleen (3 DPI, 1/4), liver (7 DPI, 1/4), bladder (7 DPI 1/4), and kidney (7 DPI, 1/4) samples (Fig 2 E). Of note, while the same bat was HeV RNA positive for both the bladder and kidney samples, no infectious virus could be isolated. No viral RNA was detected in any of the tissues collected from NiV-M inoculated bats at either 3- or 7 DPI (Fig 2E).

**Fig 2.**
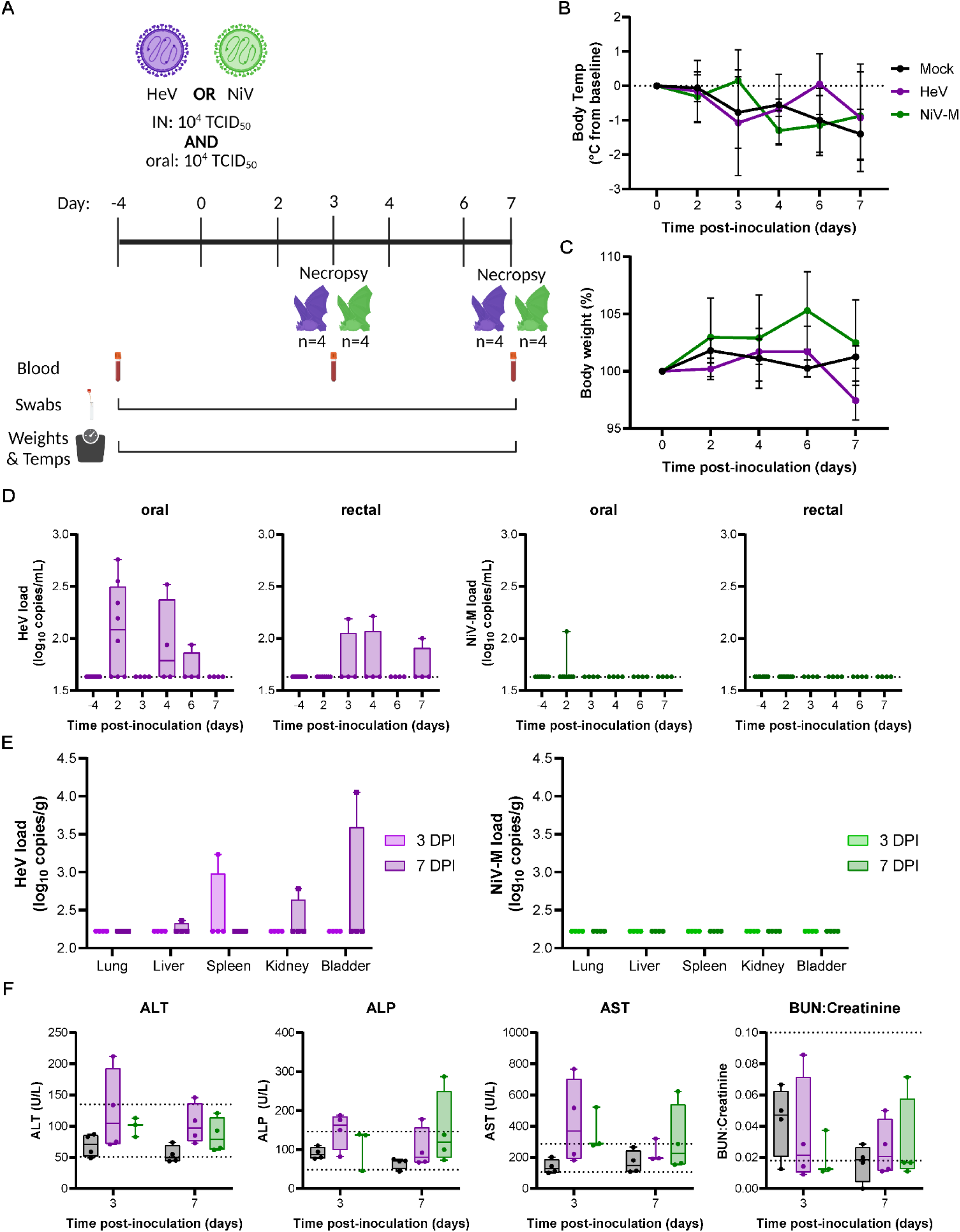
Jamaican fruit bats rapidly control Hendra and Nipah virus after intranasal and oral inoculation without signs of disease. Jamaican fruit bats inoculated with Hendra virus (HeV) (purple) or Nipah virus Malaysia (NiV) (green) via intranasal (10^4^ Tissue culture infection dose 50 (TCID)/mL), and oral (10^4^ TCID_50_/mL) routes. (A) Depiction of study (Image created in BioRender). Change in body temperature (B) and percent body weight (C) of mock bats (n=4) or bats inoculated with HeV (n=4) or NiV (n=4) followed through day 7. RT-qPCR for HeV or NiV RNA in the oral or rectal swabs (D) or tissues (E). Limit of detection 1.63 log10 copies/mL (D) or /g (E). (F) Serum levels of alanine aminotransferase (ALT), aspartate aminotransferase (AST), alkaline phosphatase (ALP) or blood urea nitrogen (BUN) to creatine ratio. Dotted lines indicate upper and lower limit in healthy control Jamaican fruit bats (n=20). Data plotted as mean ± S.D (A, B) or box defines the upper (75th percentile) and lower (25th percentile) quartiles with whiskers extending from minimum to maximum with all values shown, and the line as the median (D-F). Differences in body temperature, body weight, and serum enzyme and metabolic parameters for inoculated bats were compared to mock bats using a two-way ANOVA with Dunnett’s multiple test comparison. *P* ˃ 0.05 is not shown.

Following inoculation with HeV or NiV-M, no significant gross or histopathologic lesions were observed, consistent with the low level or lack of viral replication. Serum liver enzymes, ALT, ALP, and ASP, and BUN-to-creatinine ratio of HeV- or NiV-M inoculated bats did not differ significantly from the mock group, despite trending elevations of ALP and AST (Fig 2F).

Overall, these results suggest that JFBs can support low levels of HNV replication without overt clinical signs, but infection is controlled rapidly. Since only one oral swab was NiV-M positive, HeV was chosen for a follow-up study to evaluate if susceptibility was infection route dependent.

### Jamaican fruit bats are susceptible to Hendra virus after intravenous administration and shed into the environment

To bypass mucosal barriers to disseminated infection, JFBs were inoculated IV with 1.0 x 10^6^ TCID_50_/mL of HeV to evaluate susceptibility (Fig 3A). None of the HeV IV inoculated bats had significant change in body temperature (Fig 3B) or weight (Fig 3C) compared to mock-inoculated bats. Oral swabs were robustly positive for HeV RNA at 3 (1/4), 6 (4/4), and 7 (3/4) DPI with a peak at 6 DPI (average 4.1 log_10_ copies/mL) (Fig 3D). The viral RNA positivity of rectal swabs was more sporadic with one or two samples from different bats testing positive from 3 to 7 DPI (Fig 3D). Catch pan swabs were also collected to evaluate the potential for shedding into the environment. A pool of urine and feces swabbed in the catch pan from the male cage was HeV RNA positive on 6 DPI, supporting that HeV RNA could be shed into the environment (Fig 3D). Infectious virus was isolated from a single oral swab collected on 6 DPI, but swab infectious titers were below the limit of quantification. Most tissues and blood samples evaluated on either 3 or 7 DPI were positive for HeV RNA (Fig 3E). Viral RNA levels were high particularly in the kidney (average 3 DPI: 7.4 log_10_ copies/g, 7 DPI: 8.0 log_10_ copies/g) and spleen (average 3 DPI: 8.3 log_10_ copies/g, 7 DPI: 7.8 log_10_ copies/g). Infectious HeV was isolated from one kidney sample collected on 7 DPI, but infectious virus was below the limit of quantification for all tissue samples. Serum liver enzymes, ALT, ALP, and ASP, and BUN-to-creatinine ratio of HeV IV inoculated bats did not differ significantly from mocks, despite trending elevations of AST at 3 DPI (F 3F).

**Fig 3.**
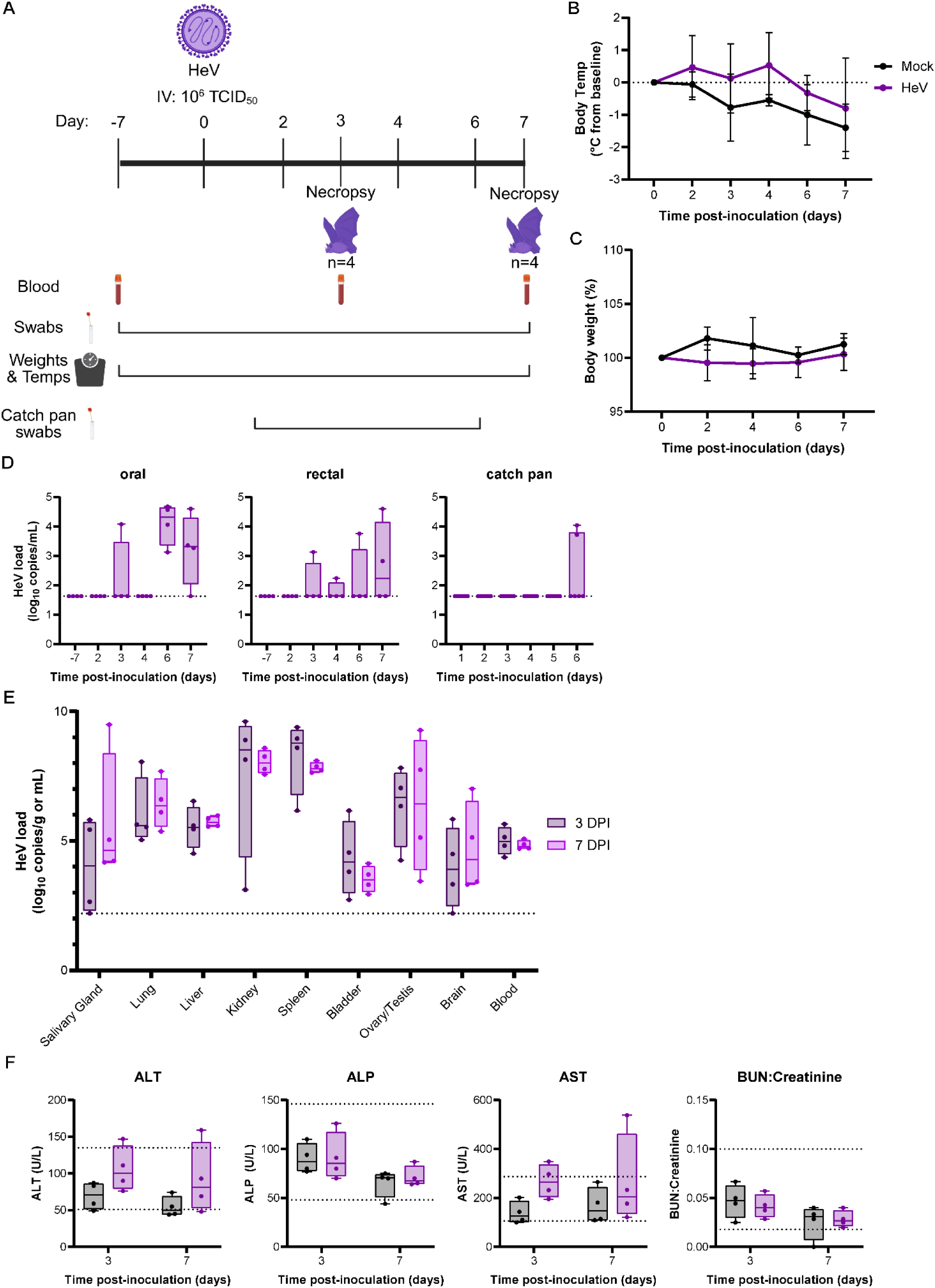
Jamaican bats are susceptible to Hendra virus and shed into the environment. Jamaican fruit bats were inoculated with Hendra virus (HeV) (purple) via the intravenous (IV) (10^6^ Tissue culture infection dose 50 (TCID)/mL) route. (A) Depiction of study (Image created in BioRender). Change in body temperature (B) and percent body weight (C) of mock bats (n=4) or bats inoculated with HeV (n=4) followed through day 7. HeV viral load in oral or rectal swabs (D) or tissues and blood (E). Limit of detection 1.63 log10 copies/mL (D) or /g (E). (F) Serum levels of alanine aminotransferase (ALT), aspartate aminotransferase (AST), alkaline phosphatase (ALP) or blood urea nitrogen (BUN) to creatine ratio. Dotted lines indicate upper and lower limit in healthy control Jamaican fruit bats (n=20). Data plotted as mean ± S.D (A, B) or box defines the upper (75th percentile) and lower (25th percentile) quartiles with whiskers extending from minimum to maximum with all values shown, and the line as the median (D-F). Differences in body temperature, body weight, and serum enzyme and metabolic parameters for inoculated bats were compared to mock bats using a two-way ANOVA with Dunnett’s multiple test comparison. *P* ˃ 0.05 is not shown.

Histopathologic evaluation of collected tissues was largely unremarkable. Minor histopathologic changes were noted focally in one section of lung in a singular IV HeV inoculated bat (S Fig 1). The lesion included focal infiltration of mononuclear cells with accumulation of foamy macrophages in adjacent alveoli and reactive endothelial cells in adjacent small caliber blood vessels. Histopathologic changes were associated with presence of viral antigen in endothelial cells and leukocytes by immunohistochemistry (S Fig 1). No significant histopathologic lesions were observed in evaluated sections of brain, cervical lymph node, salivary gland, tonsil, thymus, liver, kidney, heart, liver, spleen, adrenal gland, urinary bladder, gonad or reproductive tract. In contrast, immunohistochemistry (IHC) revealed viral antigen in the glomeruli of 50% of IV inoculated bats and within the red pulp of 50% of evaluated spleens in the absence of histopathologic changes (Fig 4). Viral antigen was not observed in evaluated sections of salivary gland, thymus, gonad and reproductive tract. Significant non-specific immunoreactivity was observed in peripheral ganglia and within the adrenal cortex all IV inoculated and mock inoculated bats.

**Fig 4.**
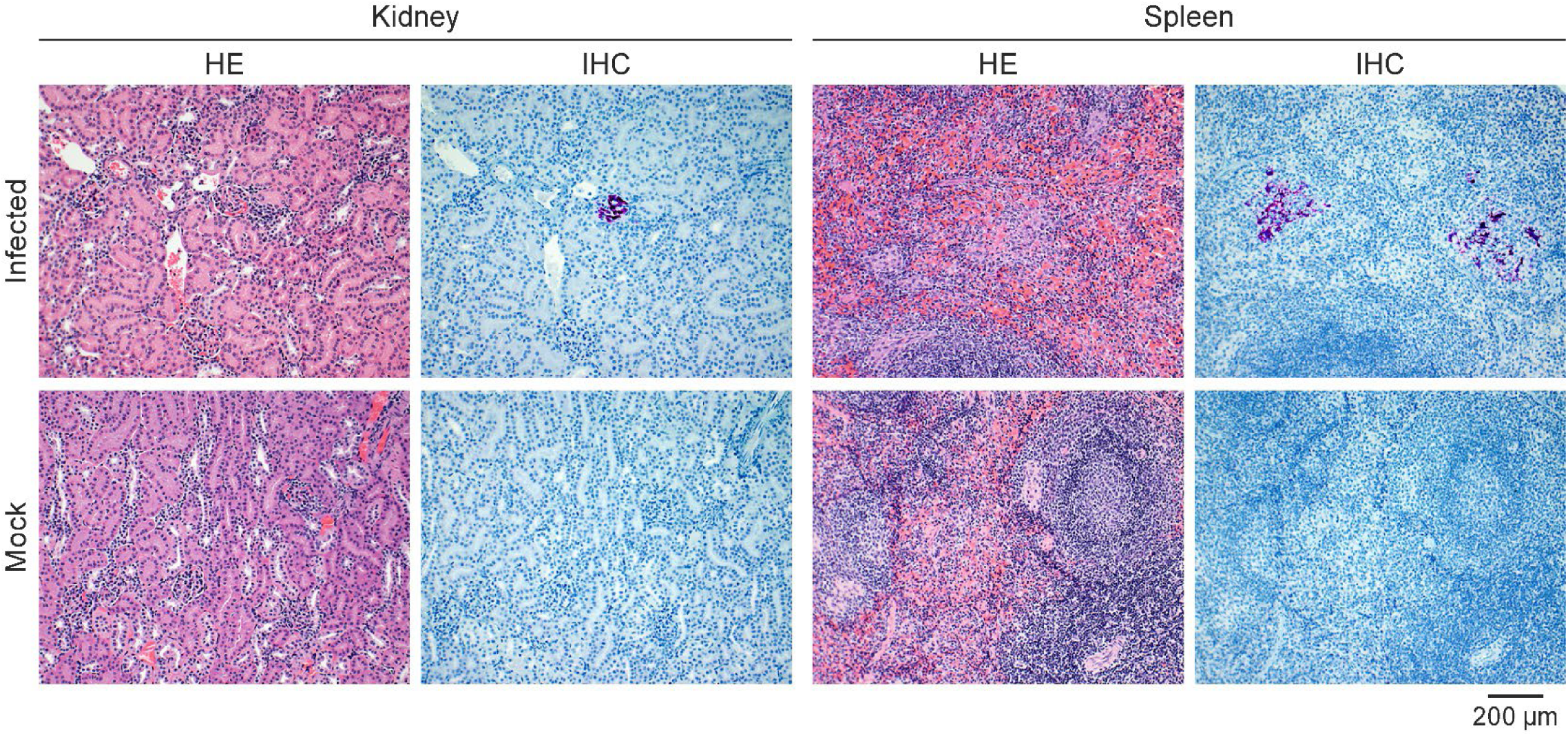
Hendra virus infection does not cause obvious histopathologic lesions in viral antigen-positive kidney or spleen. Tissue sections were stained with hematoxylin and eosin (HE) or with Nipah virus nucleoprotein antibody (purple) (IHC). Representative sections for kidney and spleen of Hendra virus intravenously inoculated or mock bats. The following magnification was used for all slides: 100X.

### Hendra virus induces mild innate antiviral responses in Jamaican fruit bats after intravenous inoculation

Host innate immune responses were evaluated following infection for both IN/PO and IV studies and compared to the mock inoculated bats. In the lung, *Ifnb1, Tnfa, Isg15, and Mx1* were significantly induced in HeV IV-inoculated bats at 3 DPI (Fig 5A). In the kidney, no *Ifnb1* transcripts were detected while *Il6* and all ISGs evaluated were significantly induced at 3 DPI with HeV IV (Fig 5B). Only *Mx1* was significantly induced in the bladder at 3 DPI in HeV IV inoculated bats, but the other ISGs trended toward induction (Fig 5C). Overall, only bats inoculated with HeV IV at 3 DPI significantly induced an innate antiviral response.

**Fig 5.**
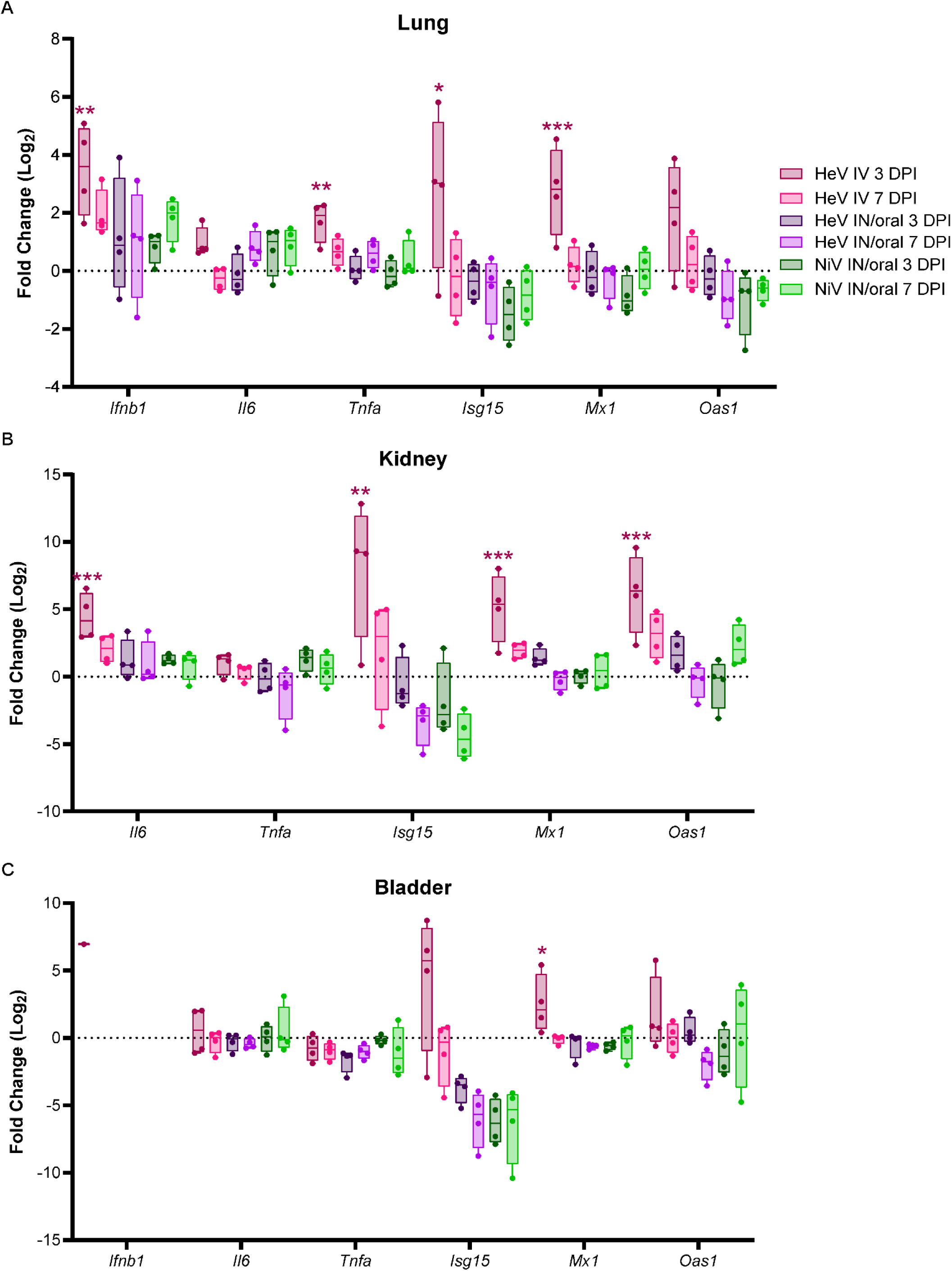
Hendra virus induces mild innate antiviral responses in Jamaican fruit bats after intravenous inoculation. RT-qPCR of interferon beta (Ifnb1), pro-inflammatory cytokine (Il6, Tnfa), or interferon stimulated gene (Isg15, Mx1, Oas1) in lung (A), kidney (B) or bladder (C) of bats inoculated with Nipah virus Malaysia (NiV) (n=4 per time point) or Hendra virus (HeV) (n=4 per time point) inoculated via the intranasal and oral (IN/oral) or intravenous (IV) routes at 3 or 7 days (d) post-inoculation. ΔC_T_ values were normalized to the average ΔC_T_ value of mock inoculated bats (n=4) to calculate ΔΔC_T_. Fold change of each gene at necropsy day was compared to the mock bats using a two-way ANOVA with Dunnett’s multiple test comparison. *P* values are indicated. **** <0.0001, *** <0.001, ** <0.01, * <0.05.

## Discussion

Although bats in the family Pteropodidae naturally host henipaviruses, experimental infections with these bats are largely impractical. A different bat model is needed to evaluate biological factors that regulate HNV susceptibility and shedding. JFBs are in the family Phyllostomidae and are distantly related to the natural HNV reservoir bat species, but they have been shown to model Ebola virus (44) and bat influenza A viruses (45, 46) effectively. Unlike pteropodid bats, JFBs are readily available to us, breed-well, and are manageable to work with in a maximum containment laboratory. Here, we demonstrated that JFBs and their cells are susceptible to infection with wild-type HNVs without immunomodulation.

JFB primary kidney cells were permissive to all HNVs evaluated. We observed robust replication of HeV, NiV-B, and NiV-M, and attenuated replication of CedV. As expected, CedV induced an innate antiviral response earlier than HeV and NiV despite replication to lower titers. Although HeV and NiV induced transcription of IFN-I, production of ISG transcripts was delayed in NiV-infected cells and only HeV infection increased ISG protein expression. Despite comparable infectious viral titers between HeV and NiV-M, IFN-I and ISG transcription remained repressed longer in NiV-M infected cells which suggests that NiV-M may antagonize the JFB innate antiviral response more effectively than HeV. The lag in innate immune activation in NiV-B compared to HeV-infected cells could be attributed to NiV-B’s slower replication.

Due to the comparable replication kinetics between HeV and NiV-M *in vitro*, these two viruses were used to evaluate JFB’s susceptibility *in vivo*. Following IN/PO inoculation, we observed limited viral RNA in oral and rectal swabs collected from each bat (n=8) inoculated with HeV. In contrast, only one oral swab was positive for NiV-M at 3 DPI. HeV RNA was detected in the kidney and urinary bladder of the same bat at 7 DPI, but no infectious virus could be isolated. Due to the prolonged attenuation of the host’s innate antiviral response in NiV-M-compared to HeV infected cells, we expected NiV-M to replicate more efficiently, but the initial IN/PO study suggested that HeV infection may be more robust. Future studies are needed to evaluate the biological determinants of viral dissemination and shedding.

Since HeV infection was more robust than NiV-M, we used HeV to evaluate Jamaican fruit bat’s susceptibility using intravenous inoculation to bypass the initial barrier of viral dissemination. In this study, all sample types evaluated, including tissues and oral, rectal, and environmental swabs, had multiple HeV RNA positive samples. Although infectious virus was below the limit of quantification, HeV was isolated from an oral swab and kidney sample collected at 6 and 7 DPI, respectively. Despite the high viral RNA loads, limited induction of innate antiviral gene transcripts was detected in the tissues. The limited innate immune responses may be due to infection being localized within the tissue preventing detection of innate antiviral responses in adjacent, uninfected regions. The bat with the minor histopathologic lung lesion at 7 DPI (S Fig 1) was not the same bat with the strongest innate immune induction in the lung (Fig 5A), which suggests either a lack of correlation between viral burden and innate responses or supports that innate responses may be localized to regions of viral antigen. Additionally, the bats could not be perfused which may have inflated the measured tissue viral loads since HeV RNA was present in the blood at necropsy (4.4-5.7log_10_ copies/mL). Future studies are needed to evaluate HNV infection in JFBs for longer periods to ensure resolution of infection, evaluate adaptive immune responses, and determine the responses that control infection in the absence of clinical disease.

The results of these infection studies in JFBs are comparable to experimentally inoculated and naturally infected flying foxes. Physiological parameters remained stable throughout infection with no changes in body weight or body temperature, no overt clinical signs, and no gross pathology (35–38). As in flying foxes, viral RNA was highest in the kidney and spleen and limited to no histopathological changes are observed near IHC positive regions (41). We also detected HeV RNA in the environment through swabbing feces and urine in catch pans immediately below their cage. This correlates with HNVs being shed in the urine of infected flying foxes (23, 25, 40).

JFB’s susceptibility to wild-type HeV *in vivo* was surprising due to their genetic distance from pteropid bats. Other studies have found that the Egyptian rousette bat (ERB) (family: Pteropodidae), a much closer relative of pteropids, does not support replication NiV *in vivo* or *in vitro* (47) despite the entry receptors, ephrin B2 and - B3, supporting entry (48). Further, ERBs do not support the replication of CedV *in vivo* and limited viral RNA was detected at 2 DPI (49). The lack of productive CedV infection in ERBs is similar to the phenotype in JFBs inoculated IN/PO with NiV-M and HeV, although at least one HeV-inoculated JFB had disseminated infection at 7 DPI, and could indicate rapid clearance rather than a lack of susceptibility. Deeper comparisons of HNV replication within *Pteropus*, JFB, and ERB *in vitro* systems could further illuminate host factors that are important in determining virus-host complementarity.

Overall, JFBs are a valuable model to evaluate bat-HNV interactions. Although the JFBs are susceptible to HeV without genetic manipulation or immune suppression, the infection is controlled rapidly comparable to HNV infection in the natural host, pteropid bats. JFB’s support of HeV replication and shedding of infectious virus enables the evaluation of the drivers of shedding and transmission. The results from future studies can be used to develop testable hypotheses that can then be addressed with free-ranging populations or captive flying foxes

## Materials and Methods

### Ethics statement

All animal experiments were conducted in an AAALAC International-accredited facility and were approved by the Rocky Mountain Laboratories (RML) Animal Care and Use Committee and adhered to the guidelines put forth in the Guide for the Care and Use of Laboratory Animals 8th edition, the Animal Welfare Act, United States Department of Agriculture and the United States Public Health Service Policy on the Humane Care and Use of Laboratory Animals. Work with infectious HeV and NiV under Biosafety level 4 (BSL4) conditions was approved by the Institutional Biosafety Committee (IBC) and conducted in RML’s BSL4 facility. For the removal of specimen from BSL4, virus inactivation of all samples was performed according to IBC-approved standard operating procedures (SOPs).

### Viruses and cells

HeV (HeV/Australia/1994/Horse) and NiV-M (NiV/Malaysia/199901924) were used for bat infection studies and in vitro experiments. CedV and NiV-B (NiV/Bangladesh/2004/human) were also used for *in vitro* studies. CedV AUS_RML_1 was isolated from a positive bat urine sample (50) as follows: Vero cells (Vero CCL-81, ATCC) in a 24-well plate were inoculated with up to 300 µL original undiluted sample and a 1:10 dilution thereof in different wells. Diluted and undiluted samples were plated in duplicates. Plates were centrifuged for 30 min at 200 × *g* and incubated for 30 min at 37 °C and 5% CO2. The inoculum was removed and replaced with 500 µl DMEM medium containing 2% FBS, 50 U/mL penicillin and 50 μg/mL streptomycin. At 2 and 3 DPI, CPE was scored. Supernatants from plates with CPE present were analyzed via RT–qPCR RNA to rule out other causes of CPE. The confirmed CedV positive supernatant was passaged once in Vero cells to generate a virus stock. All viruses were sequence verified and neither *Mycoplasma* nor contaminants were detected in any of the virus stocks. Virus was propagated in Vero CCL-81 cells in DMEM supplemented with 2% fetal bovine serum (FBS), 1 mM L-glutamine, 50 U/mL penicillin, and 50 μg/mL streptomycin (DMEM2).

#### Cells

Primary JFB kidney cells, AjKi_RML2, were maintained in DMEM/F-12 supplemented with 1X MEM non-essential amino acids (NEAA) (Gibco), 10% FBS, 50 U/mL penicillin, and 50 μg/mL streptomycin (10DMEM/F-12). Vero CCL-81 cells were maintained in DMEM supplemented with 10% FBS, 1 mM L-glutamine, 50 U/mL penicillin, and 50 μg/mL streptomycin (DMEM10). All cells were maintained at 37 °C, 5% CO2 and were *Mycoplasma* negative.

### Bat inoculations

JFBs, from a closed colony housed at Colorado State University, were shipped to RML for this study. Upon arrival at RML, bats were co-housed in same sex groups in either a flight cage, or in a stainless-steel cage with up to six bats per cage. Bats were fed fresh fruit twice daily and provided water. One week prior to the start of the study, bats were randomly assigned into same sex groups and placed in a stainless-steel cage, 4 bats per cage, to allow them to acclimate to the facility prior to inoculation. In the first study, male bats were inoculated IN (1.0 x 10^4^ TCID_50_/mL) and PO (1.0 x 10^4^ TCID_50_/mL) with HeV (n=8) or NiV-M (n=8). In the second study, equal numbers of male and female bats were inoculated IV (1.0 x 10^6^) with HeV (n=8). A group of mock bats (n=4), two male and two female, were inoculated via a combination of IN, PO, and IV routes with an equal volume of 1X Dulbecco′s phosphate buffered saline (DPBS) (IV) or DMEM (IN/PO). All inoculations and subsequent manipulations were performed under isoflurane (1–5%) anesthesia. Four HeV-, four NiV-, and two mock inoculated bats were euthanized at 3 and 7 DPI to assess viral replication and host gene expression in tissue samples. Prior to inoculation, every two days after inoculation, and at necropsy, oropharyngeal and rectal swabs were collected to monitor viral shedding. During the IV study, samples of the catch pans were collected daily approximately 6 hours after the cages were cleaned. Blood was collected at baseline and at necropsy. Whole blood was used for hematology, and serum was used for clinical chemistry. Hematology analysis was completed on a ProCyte DX (IDEXX Laboratories, Westbrook, Maine). Serum chemistries were completed on a VetScan VS2 Chemistry Analyzer (Abaxis, Union City, California). Bats’ body temperatures were obtained via implanted transponder (BMDS IPTT-300) and weights were taken on days -7, 2, 4, 6 and at necropsy. Swabs were collected in 1 mL DMEM2. Bats were observed daily for clinical signs of disease. Necropsies and tissue sampling were performed according to IBC-approved SOPs.

### Histopathology

Necropsies and tissue sampling were performed according to IBC-approved SOPs. Tissues were collected and fixed for a minimum of 7 days in 10% neutral buffered formalin with two formalin changes. Tissues were placed in cassettes and processed with a Sakura VIP-6 Tissue Tek on a 12-h automated schedule using a graded series of ethanol, xylene, and PureAffin. Prior to staining, embedded tissues were sectioned at 5 µm and dried overnight at 42 °C. Viral antigen was detected using a rabbit polyclonal anti-Nipah virus nucleoprotein antibody at a 1:1000 dilution (Cat. #GTX636902). Vector Laboratories ImmPRESS VR horse anti-rabbit IgG polymer (Cat. #MP-6401) was used as a secondary antibody. For negative controls, replicate sections from each block were deparaffinized and stained in parallel following an identical protocol, with the primary antibody replaced by Vector Laboratories rabbit IgG (Cat. #I-1000-5) at a dilution of 1:2500. The tissues were stained using the Discovery Ultra automated stainer (Ventana Medical Systems) with a Roche Tissue Diagnostics Discovery purple kit (#760-229). Histopathologic analysis was performed by a board-certified veterinary pathologist blinded to group assignments of the animals.

### *In vitro* infections

To assess henipavirus replication and host gene expression kinetics, AjKi_RML2 cells were plated at 400,000 cells/mL in 10DMEM/F-12 in 12-, 24-, or 48-well plates and incubated overnight at 37 °C, 5% CO_2_. The next day, medium was removed, and cells were inoculated with either CedV, HeV, NiV-M, or NiV-B at multiplicity of infection (MOI) 0.005 or 1.0 for 1 h at 37 °C, 5% CO_2_ with rocking every 15 min. Cells were then washed two (MOI 0.005) or three (MOI 1.0) times with DPBS, and 0.5 mL of fresh medium, DMEM/F-12 with 2% FBS, 1X NEAA, and 50 U/mL penicillin, and 50 μg/mL streptomycin (2DMEM/F-12), was added. A 300 µL sample of the medium was collected at 1 hr and every 24 hr for up to seven days and replaced with an equal volume of fresh medium. At the indicated time points, cells were lysed in 600 μL of RLT (Qiagen) with 1:100 β-mercaptoethanol for RNA extraction or 4% sodium dodecyl sulfate (SDS) buffer with 10% β-mercaptoethanol for western blotting. Protein samples were boiled at 100 °C for 10 min.

### Virus titration

Infectious virus in tissue, swab, and supernatant samples was evaluated in Vero CCL81 cells. Tissue samples were weighed, then homogenized in 1 mL of DMEM2. Vero CCL81 cells were inoculated with ten-fold serial dilutions of homogenate, swabs, blood, or supernatant incubated for 1 h at 37 °C. For tissue homogenates and blood, the first two dilutions of each sample replicate were washed twice with DMEM2. For swab and supernatant samples, cells were inoculated with ten-fold serial dilutions and the first dilution of each sample replicate was washed once. On days 5-7, cells were scored for cytopathic effect (CPE). Titers in TCID_50_/mL were calculated by the Spearman-Karber method.

### Western blot analysis

Inactivated cell lysates collected in SDS buffer were run on 4–12% Bis-Tris NuPAGE gels (Invitrogen) at 150 V for approximately 1 h then transferred onto methanol-activated PVDF membrane (BioRad). Following a 1 h block in 5% powdered milk, the membranes were washed in buffer (1X tris-HCL with 0.1% tween 20) three times. Membranes were probed with primary antibody overnight rocking at 4 °C. The next day, blots were washed three times, incubated with an HRP-conjugated secondary antibody for 1 h rocking at room temperature, and washed three times before development. Blots were incubated with a 1:1 ratio of peroxidase and enhancer reagents Clarity Western ECL (BioRad) or SuperSignal West Femto (Thermo Scientific) and developed on an iBright imaging system (Thermo Fisher Scientific). Primary antibodies used: 1:1000 pSTAT1 – Y701 (Cell Signaling Technology, 9167S), 1:1000 total STAT1 (Cell Signaling Technology, 14994S), 1:1000 total STAT2 (Cell Signaling Technology, 72604S), 1:1000 β-actin (GeneTEx, GTX629630), 1:1000 NiV N (Invitrogen, HL1436). Secondary antibodies used: 1:10,000 Donkey-anti-rabbit (GE Healthcare, NA934) and 1:10,000 Sheep-anti-mouse (GE Healthcare, NA931).

### RNA extractions and RT-qPCR

Swabs and blood were collected as described above; 140 µL was used for RNA extraction using the QIAamp Viral RNA Kit (Qiagen) according to the manufacturer’s instructions with an elution volume of 60 µL. For tissues and cells, RNA was isolated using the RNeasy Mini kit (Qiagen) according to the manufacturer’s instructions and eluted in 50 µL. Viral RNA was detected by RT-qPCR (Table 1). RNA was tested with TaqMan™ Fast Virus One-Step Master Mix (Applied Biosystems) using QuantStudio 6 or 3 Flex Real-Time PCR System (Applied Biosystems). HeV or NiV standards with known copy numbers were used to construct a standard curve and calculate copy numbers/mL or copy numbers/g.

**Table 1.**
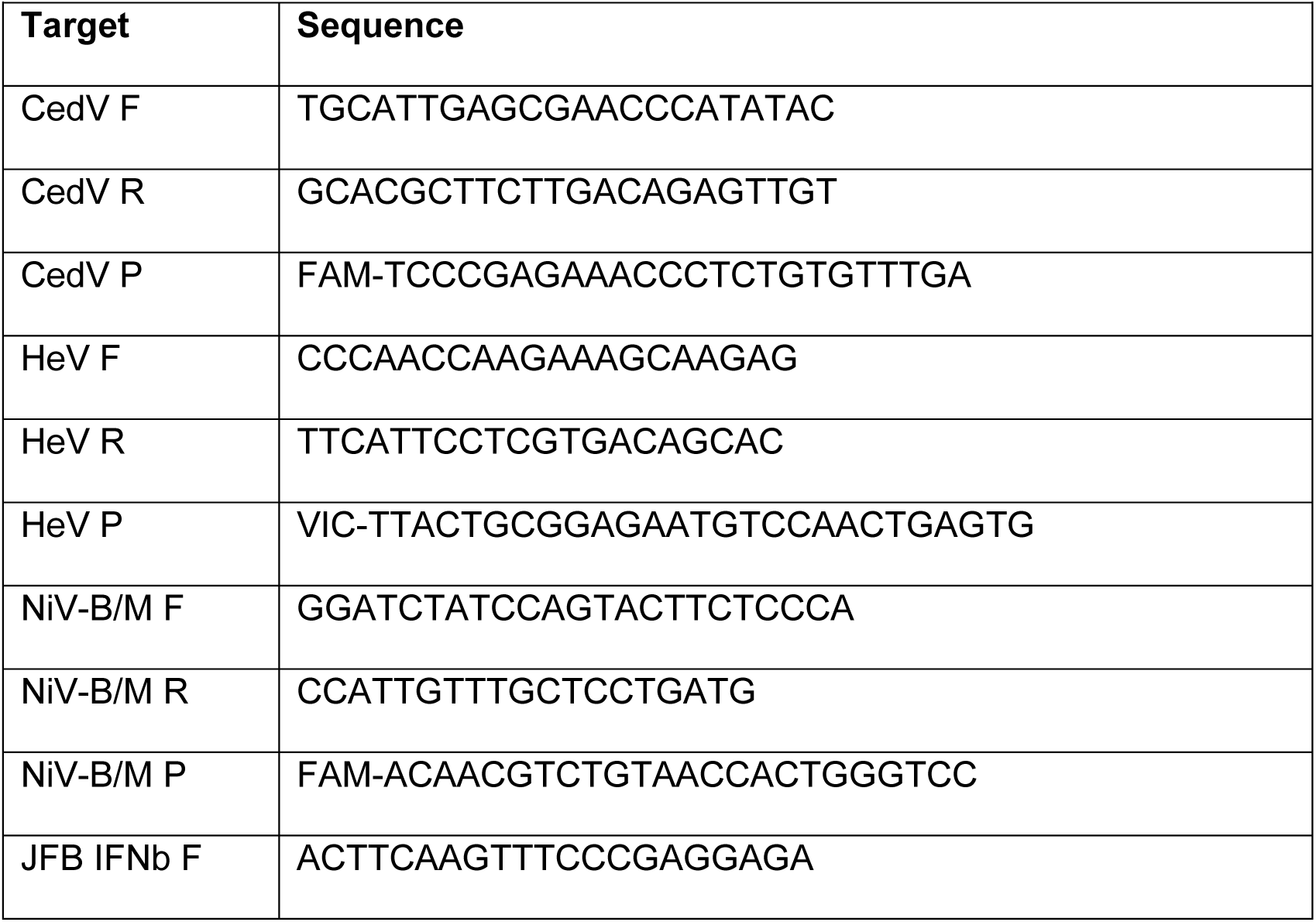
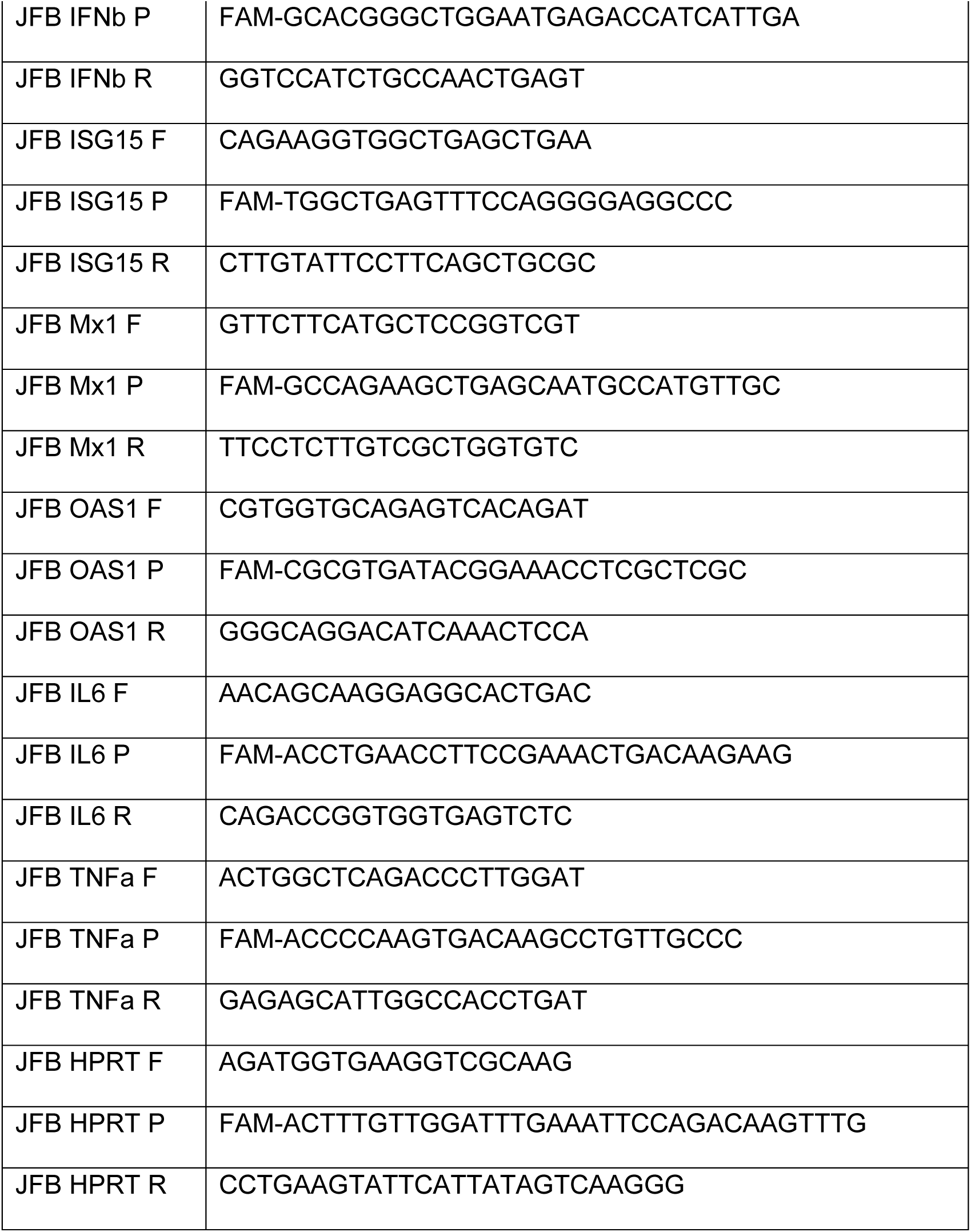
RT-qPCR primers for Jamaican fruit bat immune genes and viruses.

RLT lysates from cell monolayers were transferred to 70% ethanol prior to extraction of RNA using the RNeasy extraction kit (Qiagen). Cellular RNA was used to measure viral RNA and host genes using RT-qPCR (Table 1). Following extraction, 17 µL of RNA was treated with TURBO DNase (Invitrogen) according to the manufacturer’s instructions. After DNase treatment, the RNA was diluted 1:5 in molecular grade water (Invitrogen). DNase-treated RNA (5 µL) was used for each host gene assessed. Fold change was calculated for host genes by dividing the sample relative gene expression by the average relative gene expression of the healthy controls or mock-treated cells (2−ΔΔCt).

### Statistical analysis

Significance tests were performed as indicated where appropriate for the data using GraphPad Prism 10.2.0. Unless stated otherwise, statistical significance levels were determined as follows: no symbol = *p* > 0.05; * = *p* ≤ 0.05; ** = *p* ≤ 0.01; *** = *p* ≤ 0.001; **** = *p* ≤ 0.0001. The statistical test used is specified where appropriate.

The data generated in this study have been deposited in Figshare: https://doi.org.10.6084/m9.figshare.32310864.

## Acknowledgements

This research was supported [in part] by the Intramural Research Program of the National Institutes of Health (NIH). The contributions of the NIH author(s) are considered Works of the United States Government. The findings and conclusions presented in this paper are those of the author(s) and do not necessarily reflect the views of the NIH or the U.S. Department of Health and Human Services.

## Author Contributions

Conceptualization: SvT

Formal Analysis: SvT

Investigation: SvT, JB, BB, AW, SG, FK, JES, JRP, TB, RKM, BNW, KH, AM, JP-S, CSC

Methodology: SvT, BJS, VJM

Project Administration: SvT

Supervision: E,W, SvT

Writing – Original Draft Preparation: SvT

Writing – Review & Editing: JB, BB, AW, SG, FK, JES, JRP, TB, RKM, BNW, KH, AM, JP-S, CSC, BJS, EW, SvT

